# Translationally active soil microorganisms during substrate-induced respiration

**DOI:** 10.1101/2023.12.20.572626

**Authors:** Nina Rose Camillone, Mary Ann Victoria Bruns, Raúl Román, Daniel Wasner, Estelle Couradeau

## Abstract

Soil microorganisms carry out many processes that are fundamental to soil functions. Among the millions of microbial cells present in a gram of soil, however, less than 2% are commonly estimated to be active at any point in time. Because the respiratory response of a bulk soil to carbon substrate addition would be expected to reflect the number of active cells, we hypothesized a positive correlation between active cells and soil respiration rates during substrate-induced respiration (SIR) assays. To test this, we monitored respiration and active cell counts during 24-h incubations of agricultural soil subsamples after treating with two carbon substrates or a water-only control. We enumerated active cells with the Bioorthogonal Non-canonical Amino Acid Tagging (BONCAT) method. BONCAT provides a labeled amino acid for active cells to incorporate into newly synthesized proteins, which can then be tagged with a fluorescent dye to enable enumeration by flow cytometry. Both respiration rates and active cell counts increased over time and were positively correlated with each other after 6 h of incubation. After 24 h, increases in active cells were proportionally greater than increases in respiration. Additionally, carbon-amended soils had higher respiration rates than water-only soils with similar active cell counts, suggesting differences in carbon use efficiency. Our study documents for the first time the respiratory response from *in-situ* microbial activation induced by substrate amendment of soil within 6 h, a short enough timescale to exclude most cell replication. This study also demonstrates that the correlation between active cell numbers and respiration is substrate-dependent.

**IMPORTANCE:** While many critical ecosystem services provided by soil are known to rely on microbial activity, the soil microbial community largely remains a black box. While respiration is a common indicator of bulk soil microbial activity, this study demonstrates that the relationship between respiration and the number of active cells differs based on available carbon substrates. Advancing knowledge in this area will both enable better interpretation of biological soil tests by land managers and inform researchers modeling contributions of soil microbial respiration to global carbon dynamics.

## INTRODUCTION

Soil ecosystem services are intrinsically dependent on microbial activity through the many vital processes carried out by soil microbes. These include organic matter decomposition, plant nutrient provisioning, soil aggregate formation, nitrogen fixation, antibiotic production, and more[1], with global implications for nutrient cycling and climate change. Eighty percent of the 3200 Pg carbon in terrestrial ecosystems is stored in soil, with a flux of around 60 Pg per year due to microbial decomposition[2]. Microbial carbon use efficiency (CUE) is considered low when biomass production is accompanied by high carbon mineralization as atmospheric CO_2_, whereas high-CUE indicates more soil carbon retention in biomass per unit respiration. Variations in CUE significantly impact global carbon cycling[3].

The relationship between the soil microbiome and macroscale nutrient cycles is complicated by the soil’s high percentage of cells that are resting, inactive, or dormant[4, 5]. Since soil microenvironments are typically low in carbon and energy sources, many microbes remain dormant to survive and reactivate pending resource availability[6]. Early studies enumerating redox-active soil bacterial cells report active fractions of 5-20% or as low as 1%[7, 4, 8, 9]. More recently, however, other methods have found over 40%[10], 50%[11], or 90%[12] active bacteria in soil samples. In a laboratory incubation study, transcriptionally active soil bacterial fraction increased from near zero to 15% after 48 h[13]. This highlights that cell activity status is dynamic, with a potentially activatable fraction that may be difficult to classify in a dormant-active binary. Going forward, we define active microbes as those synthesizing new proteins and dormant microbes as those with no detectable translation to focus on potential for metabolic activity while excluding residual extracellular enzymes.

In light of the numerous beneficial functions of soil microbes, increased soil microbial activity is often considered desirable overall. For example, bulk soil respiration is a common proxy for potential microbial activity used as an indicator of agricultural soil health [14, 15]. Additionally, multiple substrate-induced respiration (MSIR) assays, such as MicroResp^©^, measure soil microbial responses to diverse C substrates, so that a broader variety of respired substrates indicates a more active and dynamic community [16, 17]. It is well established that carbon amendments initially stimulate soil respiration to a greater extent than cell division[18]. Particularly within 6 h, no shifts in community composition were observed to suggest substrate-specific reproductive selection[19]. However, it remains unknown whether reactivation of dormant microbes is an important contributor to SIR. Otherwise, bulk soil respiration may be controlled by a small percentage of already-active cells through fluctuations in their cell-specific respiration. This would indicate a major limitation for whole-community microbiome analysis in understanding and predicting C cycling. Further, experimentally and conceptually decoupling bulk community activity from the number of active microbes allows differentiation of active, efficient community members from and less efficient community members having similar respiration rates.

In this study, we investigate whether the SIR response is linked to increases in active cells relative to the total number of cells in the soil. Specifically, we employed bioorthogonal non-canonical amino acid tagging (BONCAT) and flow cytometry to enumerate active cells in tandem with measuring SIR. BONCAT enables the detection of translationally active microbes through the incorporation of a modified amino acid that is later tagged with a fluorescent dye[13, 20]. We considered that the relationship between bulk respiration and number of active cells may be dependent on the simplicity of available C substrates. Thus, as in MSIR, we used two carbon substrate treatments (glucose and galactose) to induce different respiration rates along with a water-only treatment. We measured activity after four incubation times up to 24 h. Having simultaneously monitored soil microbial activity at the bulk and cell levels, we calculated respiration rate per active cell. We hypothesized that variation in respiration rate among substrate treatments and across time would reflect changes in the number of active cells and not the number of total cells, suggesting shifts in cellular activity status. This finding would support the consideration of the fluidity of microbial activity status as a factor in modeling and managing soil ecosystem processes. Further, if bursts in respiration are strongly linked with dormant cell activation, respiration tests can be developed as proxies for the potentially active microbial fraction, facilitating future collection and interpretation of soil microbial data.

## MATERIALS AND METHODS

### Soil collection and experimental design

All soil samples were collected at the Pennsylvania State University’s Russell E. Larson Agricultural Research Center near Pennsylvania Furnace, PA (40.72°N, −77.92°W), from a long-term tillage experiment established in 1978[21]. A composite sample of bulk soil was collected in May 2023 from the top 10 cm of a long-term no-till plot planted with wheat, representing the Ap horizon of a Hagerstown silt loam (fine, mixed, semiactive, mesic Typic Hapludalfs in USDA soil taxonomy). The soil was sieved to 2 mm and stored at 4°C for less than 4 months before experimentation. Gravimetric water content was determined to be 16.6% by oven-drying a subsample at 105°C for 24 h. Available water holding capacity was determined to be 62.2% by the funnel method[42]. Incubations of soil subsamples with cycloheximide to inhibit fungi or chloramphenicol to inhibit bacteria for 6 h showed that respiration was around 60% attributable to bacteria and 40% to fungi (Supplemental Table 1)[22].

The experimental timeline is summarized in Figure **1**. Each experimental replicate consisted of 1.2 g field-moist soil (1 g oven-dried soil equivalent). Replicates were placed into 12-mL glass tubes sealable via screw-on caps with silicone septa (Exetainer®12ml Vial - Flat Bottom 738W, Labco Limited, High Wycombe, UK). The septa also allowed needle syringe access for headspace sampling. For substrate treatments, 100 µL of liquid was added to each tube: either an aqueous substrate solution containing 3.0 mol L^−1^ carbon (in the form of glucose or galactose, Sigma-Aldrich, Saint Louis, MO, USA) or ultrapure water. This resulted in a final concentration equivalent to 30 mg glucose g^−1^ soil water, which is the concentration commonly used with the MicroResp*^TM^* system[16, 17]. The gravimetric water content of the soil after addition of the substrate treatments was 30%, which was 50% of the available water holding capacity. The tubes were sealed to limit moisture evaporation and incubated in the dark at 22°C for 2, 6, 12, or 24 h. The time intervals reflect previous studies where respiration rates in surface horizon soil microcosms amended with glucose peaked from 5-10 h (depending on glucose concentration) and returned to near initial rates after 24 h[23]. Galactose was selected as a substrate treatment because it stimulates a soil respiration rate intermediate to those of glucose and water-only[16]. Initial pilot experiments included lignin and vanillic acid as additional substrates, but these treatments resulted in very low BONCAT activity levels (on average less than 0.5%) compared to the false positive discovery rate (0.2%) even after 24 h incubation. These substrates were not included, and further research will be necessary to understand why the BONCAT signal was so low in these treatments. Six replicate tubes were included per substrate treatment per incubation length.

**FIG 1.**
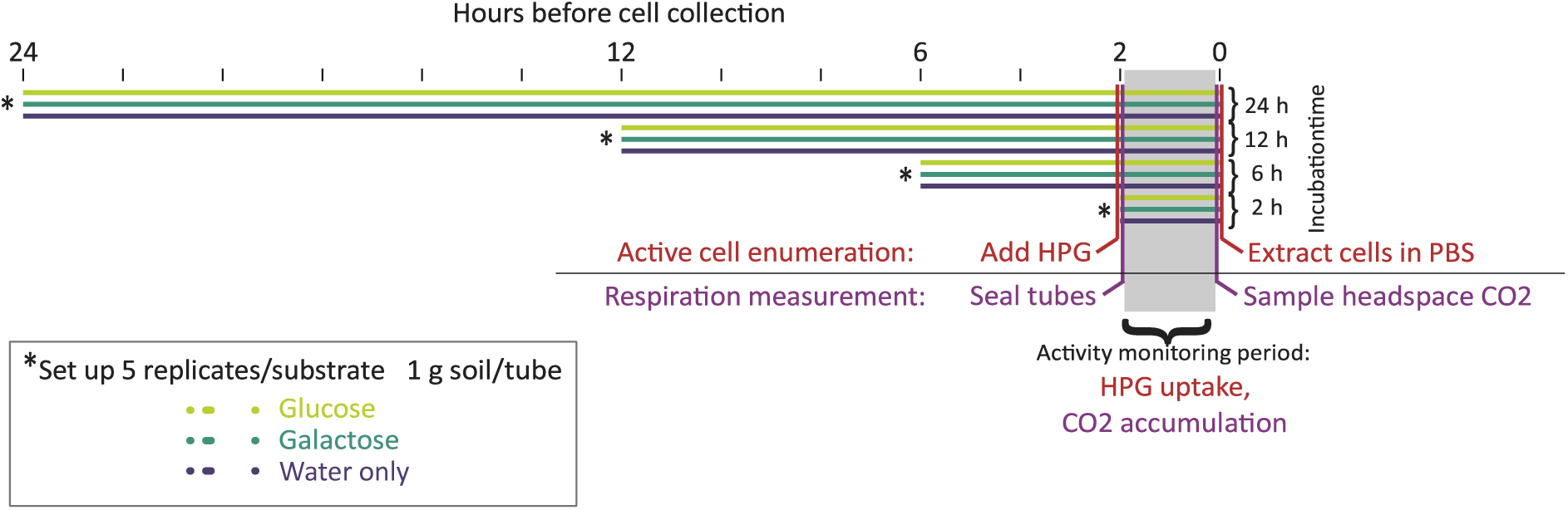
Experimental design as a timeline of substrate incubations, headspace sampling, and cell collection. L-homopropargylglycine (HPG) is a synthetic amino acid homologous to methionine used to label translationally active cells in the BONCAT workflow.

### Activity monitoring: respiration measurement and cell extraction

Two hours prior to the end of the incubation period (i.e. concurrently with the start of the substrate treatment for the 2-h replicates), the tubes were opened, flushed with ambient air, and resealed to begin a CO_2_ accumulation period of equal length for all replicates. While the tubes were open, 250 µL of 800 µmol L^−1^ L-homopropargylglycine (HPG), a synthetic amino acid homologous to methionine was also added to 5 replicates per substrate treatment per incubation length. The sixth replicate in each group received 250 µL of ultrapure water as a control for the BONCAT dye signal. As a result, the moisture content of all replicates reached 90% of available water holding capacity. The addition of HPG initiated the first stage of the BONCAT active cell enumeration method, where translationally active cells incorporate HPG into their proteins such that they can later be distinguished from HPG-negative inactive cells[13].

Two hours later at the conclusion of the incubation period, 2 mL of headspace air was collected with a needle syringe from each tube and immediately analyzed for CO_2_ content using a LI-7000 CO_2_/H_2_O Gas Analyzer (LI-COR). A calibration curve based on standards with known CO_2_ concentration was used to determine µmol CO_2_ in experimental samples. The CO_2_ content of the ambient air was recorded and later subtracted from the headspace concentrations to calculate the soil respiration rate. Immediately after headspace sampling, 5 mL 0.02% Tween 20 in phosphate buffer saline (PBS) was added to each replicate, and tubes were shaken horizontally for 5 min at high speed to extract cells from the soil. At this point, a killed soil replicate was added to the workflow. This subsample had been autoclaved three times at 1-day intervals (15 min, 121°C) in order to provide a control for staining with no cells. After shaking, the tubes were centrifuged for 5 min at 30 x g to separate soil particles without pelleting cells. The supernatant cell suspensions were stored at −20°C in 1 mL aliquots to which 500 µL 30% glycerol was added (final concentration 10% glycerol) and were processed within one week. Three replicates (one each from 6-h glucose, 12-h glucose, and 6-h water) had to be removed from analysis because respiration rates could not be quantified due to leaky tube seals.

### Active microbial cell labeling

The methods for collecting cells and BONCAT staining with a click chemistry reaction were adapted from previously published methods [13, 24]. After thawing and briefly vortexing the frozen cell suspensions, they were centrifuged at 50 x g for 5 min to pellet any larger remaining soil particles. Previous studies have used much higher centrifuge speeds while maintaining high cell recovery[25], but for this soil we had very low cell yields when higher speeds were used to pellet soil particles, perhaps due to cell clumping. The supernatant was recovered and passed through a 35 µm filter. Then, cells were pelleted from the suspension by centrifuging at 16,000 x g for 5 min. The supernatant was poured off and discarded, and the cells were resuspended in 70.4 µL 1x PBS in 2-mL snap-cap tubes by briefly vortexing. A dye mixture was prepared and added to each tube for a final reaction concentration of 5 µmol L^−1^ amino guanidine hydrochloride, 5 µmol L^−1^ sodium L-ascorbate, 100 µmol L^−1^ copper sulfate pentahydrate (ThermoFisher Scientific, Eugene, OR, USA), 500 µmol L^−1^ tris-hydroxypropyltriazolylmethylamine (THPTA), and 5 µmol L^−1^ FAM picolyl-azide dye in a final volume of 80 µL. Unless otherwise noted, all BONCAT reagents including HPG were purchased from Click Chemistry Tools (Scottsdale, AZ, USA). The tubes were incubated in the dark for 60 min to permit the click reaction between the azide dye and the HPG incorporated into the proteins of translationally active cells. Next, samples were washed to remove excess dye by centrifuging at 16,000 x g for 5 min, resuspending in 1 mL 1x PBS by brief vortexing, and repeating these steps twice more. The resulting cell suspension was diluted in 1x PBS and passed through a 35 µm filter (Falcon Round-Bottom Tubes with Cell Strainer Cap, 5 mL). SYTO™ 59 Red Fluorescent Nucleic Acid Stain (ThermoFisher Scientific, Invitrogen, Eugene, OR, USA) was added as a counterstain to distinguish cells from clay particles. Cell suspensions were incubated in the dark for 15 min.

### Flow cytometry

Cell enumeration was performed using the LSR Fortessa flow cytometer (BD Biosciences, Franklin Lakes, NJ, USA) at the Pennsylvania State University Flow Cytometry Core, which is equipped with 405 nm (violet), 488 nm, (blue), 532 nm (green), and 640 nm (red) lasers. The BONCAT dye, FAM-picolyl azide (490 nm excitation, 510 nm emission), was captured in a green channel (“FITC”) off the 488 nm blue laser. The DNA counterstain, SYTO59 (622 nm excitation, 645 nm emission), was captured in a red channel (“APC”) off the 640 nm red laser. Forward and side light scatter were also collected for all samples. All samples were run on the low flow rate setting (which we measured to be 22.6 µL/min for our samples). Data was collected for 120 s. Knowing the flow rate of the instrument enabled calculation of the original concentration of cells in the soil using the number of cells detected, the volume of extract processed, and the mass of soil extracted.

As a negative control, a “cell extract” from a soil sample that had been autoclaved 3 times at 1-day intervals (killed control) was counterstained with SYTO59 to set the threshold for distinguishing cells from soil mineral particles. Similarly, soil samples incubated with water instead of HPG solution (HPG-negative control) were used to set the threshold for BONCAT positive cells, since they should not retain any of the BONCAT dye.

Data analysis was performed using FlowJo Software (Tree Star, Ashland, OR, USA). First, forward and side scatter fluorescence thresholds were set to constrain the size of particles examined. Secondly, a threshold for SYTO positive (SYTO+) particles, which are assumed to be cells rather than clay particles, was determined based on fluorescence intensity in the APC channel from the killed (autoclaved) control samples counterstained with SYTO59. This step accounts for any potential autofluorescence from or staining of soil particles. Finally, a threshold for BONCAT positive (BONCAT+) cells was determined based on the fluorescence intensity in the FITC channel from the SYTO+ population (cells) in the HPG-negative control. To account for electronic noise from the flow cytometer or any potential retention of dyes on soil particles, the number of SYTO+ events detected in the killed control sample was subtracted from the SYTO+ count in other samples within each batch. Additionally, the number of BONCAT+ events detected in the HPG-negative control was subtracted from the BONCAT+ population in experimental samples. Any sample where the number of BONCAT+ events was less than or equal to the number of BONCAT+ events in the counterstain-only control was recorded to have zero active cells. Preliminary work using the LIVE/DEAD™ *Bac*Light™ Bacterial Viability and Counting Kit, for flow cytometry (ThermoFisher Scientific, Invitrogen, Eugene, OR, USA) detected negligible percentages of dead microbes recovered using our extraction method (<0.65% on average; see Supplemental Table 2), likely due to our cell extraction method failing to preserve cells without intact membranes, i.e. those deemed dead by the staining kit used. Thus, we report total cells as representative of the living community.

### Statistical analysis

Statistical analysis was conducted using R version 4.2.1[26], and figures were generated using R packages ggplot2[27] and ggbreak[28]. Substrate treatments within each incubation length and incubation lengths within each substrate were compared using one-way ANOVAs. Incubation length was treated as a categorical variable due to the non-linear relationship of response variables and time. Tukey HSD post-hoc tests were performed for variables with significant ANOVAs (p<0.05). The rate of change of numbers of active cells in each substrate treatment was calculated per unit time as if it were a growth rate, but we do not refer to this as a “growth rate” since new active cells could be the result either of cell division or activation of dormant cells. Instead, this variable will be referred to simply as the rate of change of numbers of active cells. The cell-specific respiration rate was calculated by dividing the bulk respiration rate by the number of active cells in the same experimental replicate as an indicator of resource-use efficiency. The relationship between respiration rate and log-transformed number of active cells was analyzed using linear regression.

## RESULTS

### Cell counts and bulk respiration

The total number of microbial cells ranged from 3.1 × 10^5^ to 2.2 × 10^7^ g^−1^ soil throughout the experiment (Figure **2**A). The number of active cells ranged from 1.7 × 10^3^ to 4.4 × 10^6^ g^−1^ soil (bars in Figure **2**B, left y-axis), or 0.1% to 27% of the total (percentages noted below data bars in Figure **2**A). Generally, the number and percentage of active cells increased with time for all treatments. For the glucose treatment, numbers of total and active cells were highest after 12 h of incubation. In galactose, total and active cells were highest at 24 h. For the water treatment, total cells were highest at 12 h while active cells were highest at 24 h. Among the treatments, the glucose treatment had the highest total number of cells throughout the experiment while the total cells in the galactose and water treatments never significantly differed from each other. As for active cells, they likewise were highest in glucose compared to the other treatments at 6 h, 12 h, and 24 h. In water, active cell numbers averaged slightly lower than in galactose at 6 h and 24 h, but they were significantly higher than in galactose at 12 h. Contrastingly, galactose had the most active cells at 2 h, although it was still low at 2% of total cells.

**FIG 2.**
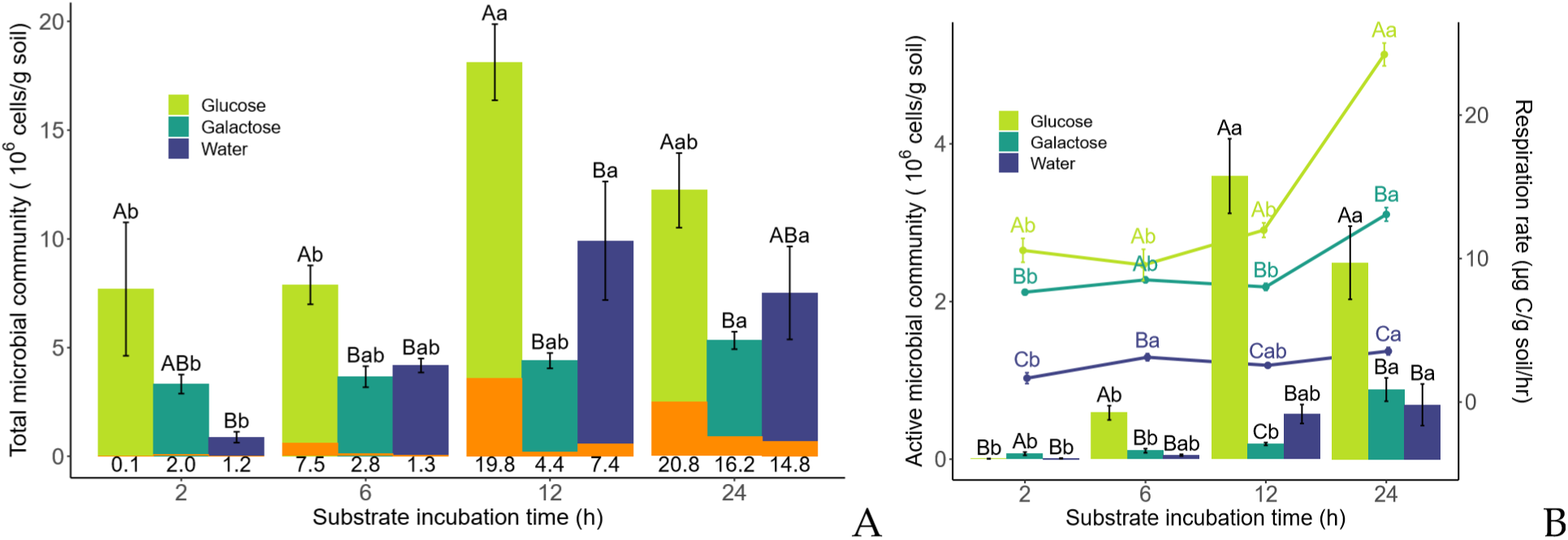
Total cells, active cells, and respiration rate. (A) Total cells over time for different substrate treatments, reported in cells per g dry soil. The lower (orange) portion of each bar represents the active cell fraction. Numbers below each bar indicate the percentage of active cells. (B) Active cells (bars, left y-axis) and respiration rate (lines, right y-axis). Note that 0 for the right y-axis is elevated. For both (A) and (B), uppercase letters represent statistically significant differences found by Tukey’s HSD test (p<0.05) among different substrate treatments within the same time point, indicated by the x axis. Lowercase letters represent statistically significant differences across time points within the same substrate treatment, indicated by color. The absence of a letter of a particular case above a bar indicates that Tukey’s test was not conducted due to an insignificant ANOVA result. Bars indicate standard error.

The bulk respiration rate increased after 24 h in all 3 treatments (lines in Figure **2**B, 267 right y-axis). The respiration rate of the substrate-amended treatments did not significantly differ from their initial rates until 24 h. Glucose consistently had a higher respiration rate than galactose, and galactose consistently had a higher respiration rate than water-only.

The largest absolute increases in numbers of total and active cells coincided with each other for all three treatments. In glucose and water, they were between 6 and 12 h, but in galactose they were later, between 12 and 24 h. For glucose, the largest increase in respiration rate occurred between 12 and 24 h, after the large increase in number of active cells between 6 and 12 h. For galactose, the largest changes in number of active cells and respiration coincided between 12 and 24 h. For water, the largest increase in respiration occurred first, between 2 and 6 h, before the increase in active cells at 12 h. Glucose had the largest changes in total and active cells and in respiration rate compared to the other treatments.

### Cell-specific respiration rate

Cell-specific respiration was calculated for each replicate by dividing bulk respiration by the number of active cells. In all three treatments, the cell-specific respiration rate was highest after 2 h and decreased thereafter. However, this decrease was only statistically significant for glucose (Figure **3**). At 2 h, the glucose treatment had the highest cell-specific respiration, driven by low active cell counts and a high respiration rate. At 12 h, cell-specific respiration was highest for galactose, caused by lower active cell counts than glucose and higher respiration rates than water. After 6 h and after 24 h, there were no statistically significant differences in cell-specific respiration rate among treatments.

**FIG 3.**
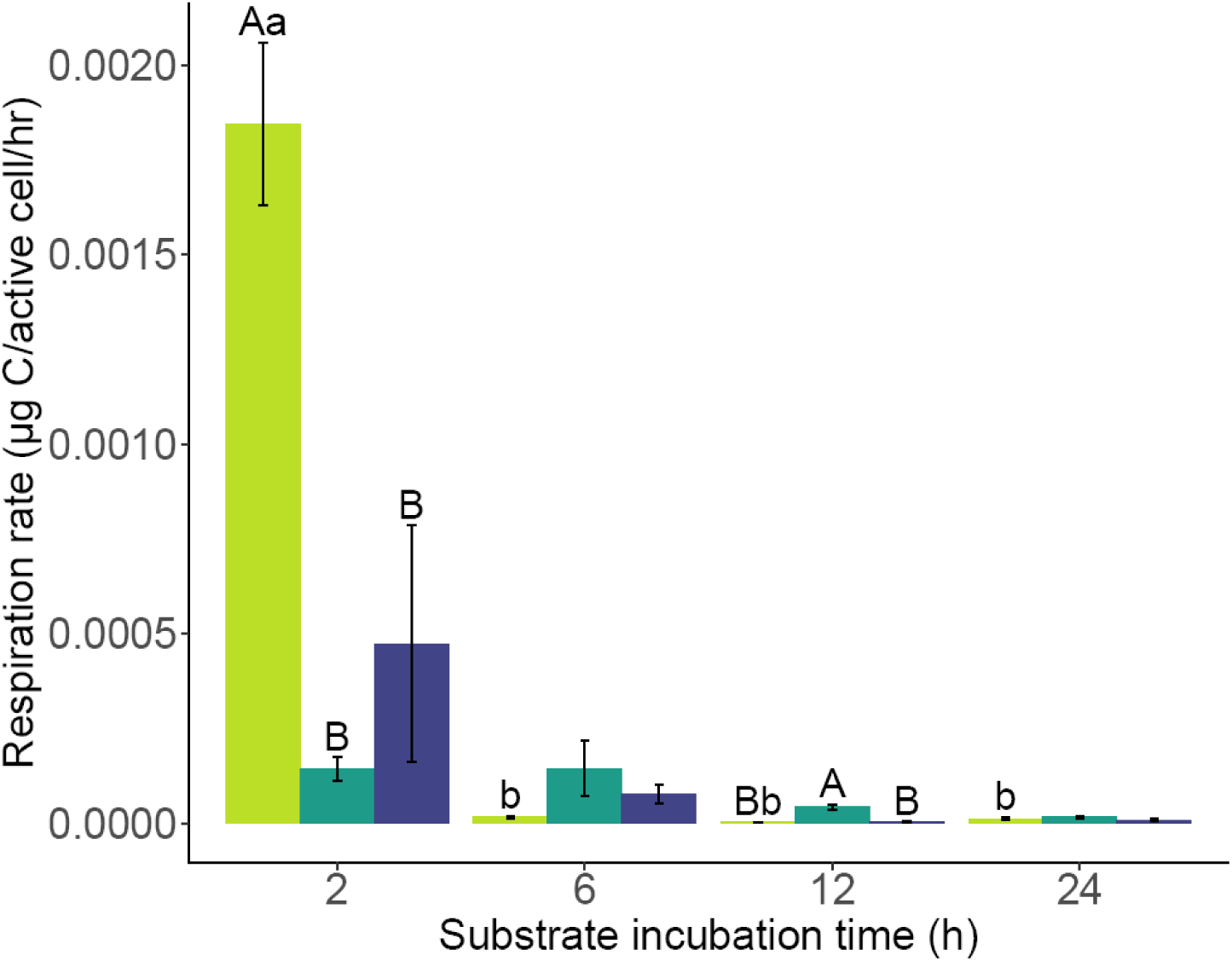
Cell-specific respiration rate, calculated by dividing bulk respiration by the number of active cells. Uppercase letters represent statistically significant differences found by Tukey’s HSD test (p<0.05) among substrate treatments within the same time point, indicated by the x axis. Lowercase letters represent statistically significant differences across time points within the same substrate treatment, indicated by color. The absence of a letter of a particular case above a bar indicates that Tukey’s test was not conducted due to an insignificant ANOVA result. Bars indicate standard error.

## DISCUSSION

### Methodological insights and comparisons

Our study documents for the first time *in-situ* soil microbial cell activation induced by substrate amendment within 6 h, a short enough timescale to exclude most cell replication. This is possible using the BONCAT method because it detects protein translation rather than cell growth, which has previously been used to monitor soil microbial activity via methods such as SIP[29]. A longer activity labeling period can increase the detection of low-activity cells but does not give a snapshot for time-resolved changes in cell activity status. Other methods for enumeration of active soil microbes involve dyeing active cells post extraction[5, 9, 10, 11], whereas in BONCAT active cells are tagged *in situ* with a synthetic amino acid prior to extraction and subsequently dyed through a click-chemistry reaction. This could lead to discrepancies between detected and actual activity if dormant cells are reawakened or if activity is suppressed through the disturbance of extraction, e.g. aerobic respiration decreasing upon saturation. Differences among soil active microbes labeling methods are summarized in Table **1**. Comparative studies are needed to determine if variation in reported numbers of active cells reflects actual cell physiology (e.g. some cells contain active enzymes but are not currently translating new ones), natural differences in soil samples across time and space, or inefficiencies in some methods compared to others.

**TABLE 1.** Comparison of methods to enumerate active microbial enumeration in soil. Abreviations: 5-cyano-2,3-ditolyl tetrazolium chloride (CTC, an artificial electron acceptor), 5(6)-carboxyfluorescein diacetate (CFDA, a substrate hydrolyzed by intracellular esterases), fluorescence *in situ* hybridization (FISH), stable-isotope probing (SIP).

| Method | CTC | CFDA | FISH | DNA-SIP | BONCAT |
| --- | --- | --- | --- | --- | --- |
| Activity type | Aerobic respiration (redox) | Metabolic enzymes | rRNA content | DNA replication | Protein translation |
| Time scale | 30+ min | 30+ min | Fixed cells | 1+ days | 30+ min |
| Labeling location | Extracted cells | Extracted cells | Extracted cells | *in situ* | *in situ* |
| Reported active fraction | 2-6%[9]<br><1%[5] | 42%[10] | 55%[11] | 91%[12] | 25%[13] |

Some remaining methodological challenges in active cell enumeration are specific to BONCAT, while others apply more broadly. For example, BONCAT does not detect the activity of microbes with access to native methionine (outcompeting analog HPG) or microbes with adequate protein reserves without need for translation. Regardless of staining method, extraction cannot capture all microbes present in the soil, especially filamentous microbes such as fungi. Typically, soils contain 10^7^-10^10^ prokaryotic cells per gram[30], compared to our counts of 5 × 10^6^ after 2 h of incubation (Figure **2**A). We confirmed that our cell extracts had very low numbers of dead microbes (Supplemental Figure 1), which may contribute to our lower cell counts considering previous research reporting 40% dead microbial cells in soil[7]. We assumed that our percentages of active cells were representative of the whole live community. In the end, any binary divide between active and inactive cells is inherently a simplification, yet it can still yield insights into overall community status for the purpose of this discussion.

### Glucose amendment reactivated dormant cells

Active cell numbers increased most rapidly from 2-6 h of incubation with glucose (Figure **4**). Notably, this was our only observation of an increase in total cells that was outpaced by a larger increase in active cells. In other words, the number of dormant cells (estimated by subtracting active from the total) decreased, suggesting cell activation. Alternatively, or in addition, this could indicate a death rate among inactive cells outpaced by reproduction among active cells; however, this seems less likely considering known soil microbial growth rates (depicted in Figure **4** as rate of change per h). Assuming no growth or reactivation of dormant cells, active cells would have been doubling every 37 min to achieve the observed increase between 2-6 h for the glucose treatment. This approaches the theoretical maximum growth rate under optimum soil conditions with as a generation time of 0.5 h[31]. For comparison, *Eschedrichia coli* has a generation time around 0.3 h *in vitro*[32]. In actuality, soil microbes have been clocked with doubling times only as low as 7.2 h in the rhizosphere[33] and 14 to 45 days in bulk soil[34] (depicted in Figure **4** as rate of change per h). This suggests that reactivation of dormant cells largely contributed to the increase in active cells. With the same reasoning of a high rate of increase in active cells, dormant microbes were likely reactivating in the water treatment between 2-12 h (Figure **4**). The rate of change in active cell counts in the galactose treatment did not measurably change throughout the experiment and remained comparable to a highly active rhizosphere environment. The growth rates for total cells were generally much lower than active cells’ (Supplemental Figure 1).

**FIG 4.**
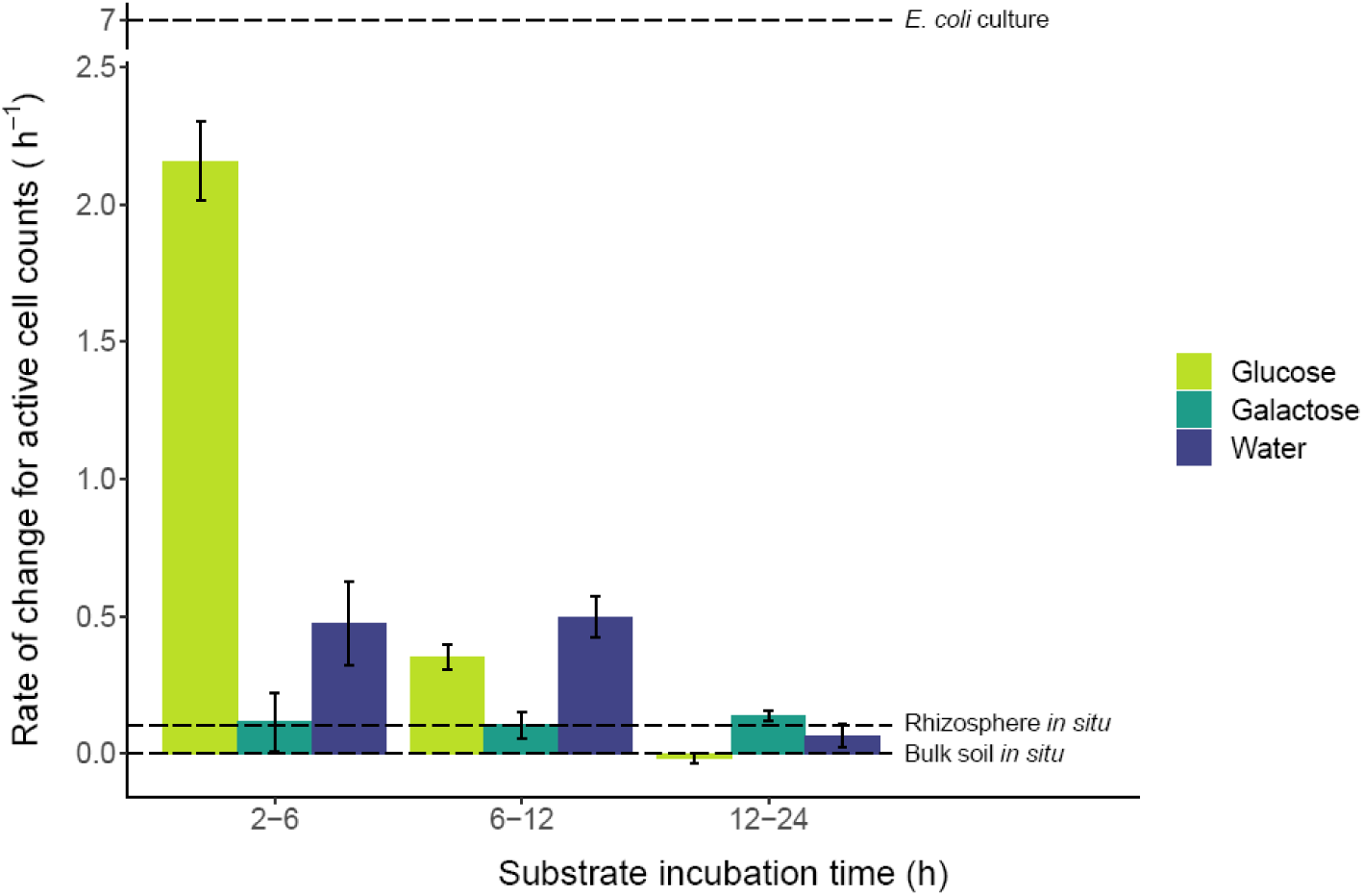
Rate of change in number of active cells over time. Bars indicate standard error, propagated from the standard error in the enumerated community sizes. For comparison, dashed lines indicate several known growth rates: a laboratory culture of *Escherichia coli* (generation time 0.3 h)[32], an active rhizosphere community (generation time 7.2 h)[33], and bulk forest soil (generation time 14.1 days) [34]. Note the break in the y axis between 2.5 and 7.

### Cell-specific activity differed by treatment

As hypothesized, respiration rates and active cell counts were positively correlated, although the relationship differed by substrate. Regression analysis revealed statistically significant relationships between these variables within each treatment and across all treatments together, although in the glucose treatment it was only significant at the p<0.06 level (equations reported in Figure **5**A). Differences among the regression models’ slopes or intercepts result in the regression line for glucose being higher than that of galactose, which is in turn higher than water’s. This pattern reflects overall trends in the treatments’ cell-specific respiration across the experiment. Lower respiration and active cell counts in the galactose treatment than the glucose treatment are consistent with the ability of cells to utilize glucose directly. By contrast, galactose requires multiple enzyme-catalyzed reactions before a downstream metabolite enters the glycolytic pathway [35].

**FIG 5.**
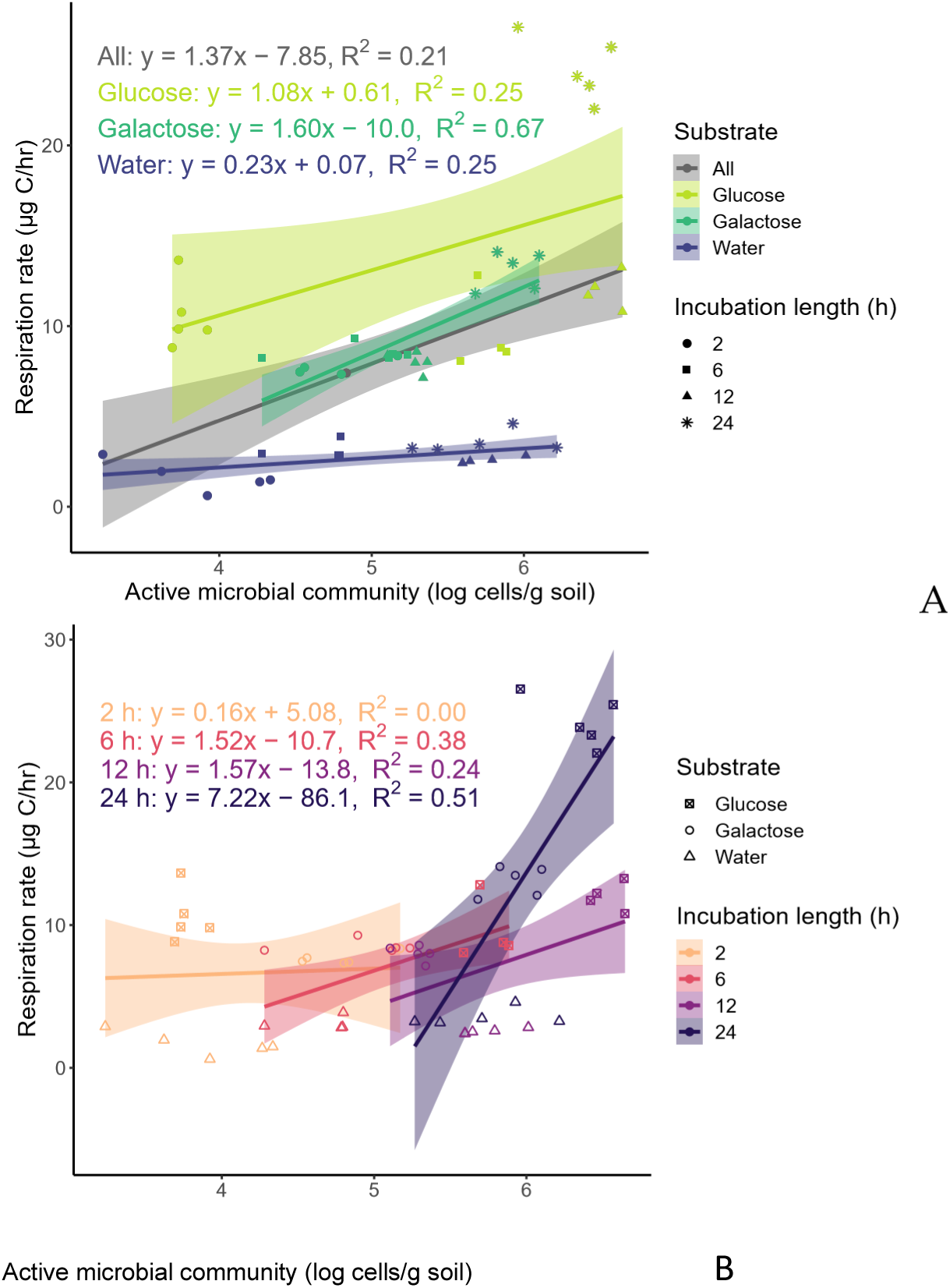
Regressions of bulk respiration onto active cell counts in log scale. The shaded regions show the 95% confidence region for each regression line. (A) Regression lines for all data and for each substrate individually. P-values for coefficients and intercepts respectively are <0.001 and 0.08 for all data, 0.06 and 0.9 for glucose, <0.0001 and 0.009 for galactose, and 0.03 and 0.9 for water. (B) Regression lines for each time point individually. P-values for coefficients and intercepts respectively are 0.9 and 0.6 for 2 h, 0.03 and 0.2 for 6 h, 0.08 and 0.2 for 12 h, and 0.003 and 0.008 for 24 h.

Without any carbon addition, the water-only treatment has the lowest regression coefficient (Figure **5**A). This indicates that this moist soil environment supported an expanding active community with minimal changes in respiration rate. This finding is consistent with the Microbial Efficiency-Matrix Stabilization (MEMS) framework, which theorizes that respiration is highest where inputs are low quality (excess ratio of C to other nutrients) and have a low capacity to be stabilized in the mineral soil matrix [36], e.g. glucose supports lower CUE than lignin. Active microbes in the water-only, carbon-poor treatment may use carbon more efficiently, i.e. prioritize its assimilation as biomass relative to mineralization. This is also consistent with previous studies that found that soil microbes metabolized complex substrates with a higher carbon use efficiency[37]. (also check decreased CUE with excess C in [38])

### Lag between respiration response and increase in active cells

Within 2 h of substrate addition, the number of active cells did not differ among treatments (Figure **2**), while the two carbon-amended treatments already had much higher bulk respiration rates than water-only (Figure **2**B). Thus, our results indicate an initial lag where changes in translational activity are decoupled from the respiration response. As hypothesized, we observed that active cell counts and respiration rates followed similar patterns across treatments when cell-specific respiration was consistent, i.e. after 6 and 24 h of substrate incubation (Figure **3**). Regression models calculated within each time point also indicated positive correlations between respiration and active cell counts with statistically significant regression coefficients at both 6 and 24 h, but not at 2 or 12 h (Figure **5**B). At these time points, by contrast cell-specific respiration differed among treatments, reflecting weak or no correlation between respiration and number of active cells. The glucose treatment had the highest cell-specific respiration at 2 h, and galactose had the highest at 12 h (Figure **3**). These differences suggest that glucose-amended soil microbes began rapidly metabolizing with a burst of respiration per cell within 2 h before the activation of dormant cells by 6 h. Meanwhile, galactose induced increases in respiration and active cell counts at a slower but steadier rate. In fact, the galactose treatment exhibited the least variation in cell-specific respiration and the strongest positive correlation (highest regression R^2^) between bulk respiration and active cell counts (Figure **5**A). A low level of respiration from BONCAT-negative cells also would have contributed to low cell-specific respiration and a strong correlation between respiration and active cells. This could be the case if few cells already contained the enzymes necessary for galactose metabolism. Regardless, for all substrate treatments, cell-specific respiration tended to decrease over time, suggesting an initial dip in CUE with excess carbon and a lag in cell activation post substrate addition. In the field, this could contribute to increased carbon emissions from more frequently disturbed soils.

### Implications for soil testing

Our results demonstrate the utility of SIR as a proxy for soil microbial activation in an agricultural soil. Soil testing already utilizes SIR for crop yield prediction[39, 40]. Thus, future research in more soils establishing a correlation between SIR and numbers of active microbes would also implicate a correlation with agricultural yields. This may seem intuitive, considering the microbial basis for many critical agricultural soil functions such as decomposition and aggregate formation. However, other factors complicate the relationship between microbial activity and optimal soil functionality. Frequent tillage disturbance, for example, could over-stimulate microbial activity and cause faster depletion of soil organic matter with time. Additionally, CUE can affect the relationship between respiration and microbial functionality. A high respiration rate in an SIR test can indicate a responsive microbial community or simply a less efficient one.

However, microbial researchers have described activity status not simply as a dichotomy between dead and alive or between active and dormant, but as a spectrum of activity levels, defining potentially active microbes as those that resume activity after 4-12 h of stimulation [7]. While a large active microbial fraction could achieve a high respiration rate and rapidly deplete soil carbon stores, a large *potentially* active fraction would be a better indicator of a responsive and resilient microbiome supporting soil functions. The present study found rapid increases in active cell counts but not total cell counts for 6 h of substrate incubation. Previous MSIR studies have found 6-h incubation time to be optimal for observing the existing community while limiting time for shifts in community composition[16, 17, 19], which our study corroborates. Additionally, our results showed a similar effect of substrate treatment on respiration rate and active cell counts at this time (Figure **2**B), and the earliest correlation between these two indicators (Figure **5**B). These results support the interpretation of SIR tests as measures of potentially active microbes responsive to different substrates.

At the same time, active cell numbers alone could not consistently predict respiration due to differences among substrate treatments. Both carbon substrates induced higher cell-specific respiration than the water-only treatment at different times (Figure **3**), and carbon-amended replicates had higher respiration than the water replicates with similar numbers of active cells (Figure **5**A). Previously, a field study found that tilled soils with incorporation of organic residues had higher respiration rates with lower microbial biomass compared to no-till soils[41], similarly pointing toward higher activity per cell with access to carbon.

Overall, this project demonstrates a relationship between respiration and numbers of active microbes. This provides a basis for interpreting respiration-based soil tests in microbial terms and targeting research efforts on active microbes due to their contribution to essential soil functions. Future research coupling BONCAT with cell sorting and metagenomic sequencing can reveal which microbes are activated under different soil conditions and amendments. Linking other bulk soil tests to active cell counts can further inform soil management decisions for promoting desired outcomes of microbial activity.

## Supporting information

Spplementary documents

## ACKNOWLEDGMENTS

We acknowledge that gas analysis for CO_2_ content was conducted in the lab of Jason Kaye with training and support from Brosi Bradley. We would also like to acknowledge that cell enumerations were conducted at the Huck Institutes’ Flow Cytometry Core Facility (RRID:SCR_024460) on the BD Fortessa flow cytometer with training and support from multiple staff members.

## DATA AVAILABILITY STATEMENT

Original flow cytometry data are available as FCS files at doi.org/10.26208/G8KX-GG32 through Penn State’s DataCommons.

