## Supplementary material for "Translationally active soil microorganisms during substrate-induced respiration": Spplementary documents

**Supplemental table 1.** Respiration response to selective inhibition. Compounds were added to 1 g subsamples of soil along with glucose for a 6-h incubation, following the same glucose treatment and respiration measurement protocols described in the methods section. Water was added without HPG for the final 2 h of the incubation, during which CO_2_ was collected. Inhibition is reported as the percentage decrease in respiration relative to the soil with glucose and no inhibitory compound. Increased inhibition was not observed with higher dosages of the inhibitors (data not shown). Proportions of respiration attributable to bacteria and fungi were calculated by the equations (A-B)/(A-D) and (A-C)/(A-D) respectively (Anderson & Domsch 1974), where A is the uninhibited respiration rate, B is the respiration rate with streptomycin and penicillin addition, C is the respiration rate with cycloheximide addition, and D is the respiration with both inhibitor solutions. A - [(A-B) + (A-C)] was 5% different from D, the suggested acceptable allowance for experimental errors. The higher percent inhibition seen for the combined treatment than an expected additive effect of the individual inhibition treatments suggests some occurrence of non-specific inhibition.

| **Inhibitor** | **Dosage (µg/g soil)** | **Respiration inhibition** |
| --- | --- | --- |
| Cycloheximide (antifungal) | 500 | 17% |
| Streptomycin and penicillin (antibiotics) | Penicillin: 150  Streptomycin: 250 | 20% |
| Both antifungal and antibiotics | Same as above | 32% |

**Supplemental table 2.** Percentage of live cells for select samples.

| **Substrate** | **Incubation length (h)** | **Live cell fraction (mean)** |
| --- | --- | --- |
| Water | 2 | 98.9% |
| Water | 12 | 98.9% |
| Water | 24 | 99.2% |
| Glucose | 2 | 99.1% |
| Glucose | 6 | 98.9% |
| Glucose | 12 | 98.9% |
| Glucose | 24 | 99.1% |

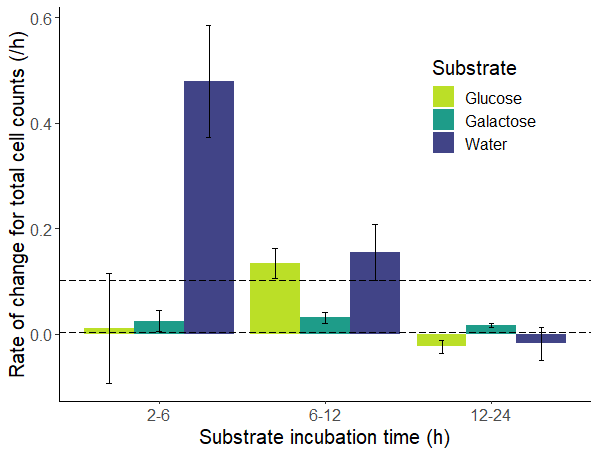

**Supplemental figure 1.** Rate of change of total cell counts, calculated as described for active cell counts in Figure 3 in the main text. The dashed horizontal lines indicate literature values for bacterial growth rates in bulk soil (lower line) and the rhizosphere (upper line).
